# A metabolic switch from OXPHOS to glycolysis is essential for cardiomyocyte proliferation in the regenerating heart

**DOI:** 10.1101/498899

**Authors:** Hessel Honkoop, Dennis E.M. de Bakker, Alla Aharonov, Fabian Kruse, Avraham Shakked, Phong Nguyen, Cecilia de Heus, Laurence Garric, Mauro J Muraro, Adam Shoffner, Federico Tessadori, Joshua C. Peterson, Wendy Noort, George Posthuma, Dominic Grün, Willem J. van der Laarse, Judith Klumperman, Richard T. Jaspers, Kenneth D. Poss, Alexander van Oudenaarden, Eldad Tzahor, Jeroen Bakkers

**Author notes:** corresponding author: Jeroen Bakkers, PhD, Hubrecht Institute, Uppsalalaaan 8, 3584 CT, Utrecht, The Netherlands. These authors contributed equally (Hessel Honkoop, Dennis E.M. de Bakker).

## Abstract

The capacity to regenerate damaged tissues, such as the heart, various enormously amongst species. While heart regeneration is generally very low in mammals ^1–3^, it can regenerate efficiently in certain amphibian and fish species ^4,5^. Zebrafish has been used extensively to study heart regeneration, resulting in the identification of proliferating cardiomyocytes that drive this process ^5–7^. However, mechanisms that drive cardiomyocyte proliferation are largely unknown. Here, using a single-cell mRNA-sequencing approach, we find a transcriptionally distinct population of dedifferentiated and proliferating cardiomyocytes in regenerating zebrafish hearts. While adult cardiomyocytes are known to rely on mitochondrial oxidative phosphorylation (OXPHOS) for energy production, these proliferating cardiomyocytes show reduced mitochondrial gene expression and decreased OXPHOS activity. Strikingly, we find that genes encoding rate-limiting enzymes of the glycolysis pathway are induced in the proliferating cardiomyocytes, and inhibiting glycolysis impairs cardiomyocyte cell cycle reentry. Mechanistically, glycolytic gene expression is induced by Nrg1/Erbb2 signaling, and this is conserved in a mouse model of enhanced regeneration. Moreover, inhibiting glycolysis in murine cardiomyocytes abrogates the mitogenic effects of Nrg1/ErbB2 signaling. Together these results reveal a conserved mechanism in which cardiomyocytes undergo metabolic reprogramming by activating glycolysis, which is essential for cell cycle reentry and heart regeneration. This could ultimately help develop therapeutic interventions that promote the regenerative capacity of the mammalian heart.

## Results

After injury, the zebrafish heart regenerates by proliferation of a small population of existing cardiomyocytes located mostly adjacent to the injury site, also known as the border zone ^6–8^. These cycling cardiomyocytes exist within a heterogeneous cell population including non-proliferating cardiomyocytes, endothelial cells and immune cells, hampering the discovery of genetic programs specific for these proliferating cardiomyocytes ^9–11^. Thus, a single cell approach is required to isolate them for detailed analysis. We generated a transgenic zebrafish *nppa* reporter line (*TgBAC(nppa:mCitrine)*) in which mCitrine expression recapitulates endogenous expression of the border zone stress gene *nppa* (which encodes a natriuretic peptide) (Extended Data Fig. S1). Histochemical analysis of injured adult hearts revealed that 75% (± 7%, n=3 hearts) of the *nppa:mCitrine* expressing cells were proliferating cardiomyocytes (Fig. 1a). To isolate these proliferating border zone cardiomyocytes, *nppa:mCitrine* hearts were cryo-injured, followed by cell dissociation and FACS sorting of mCitrine^high^ cells (Fig. 1b). In addition, we FACS sorted mCitrine^low^ cells to obtain non-proliferating remote cardiomyocytes from the same tissue for comparative analysis. Individual, living cells were sorted, followed by single-cell mRNA-sequencing using the SORT-seq (SOrting and Robot-assisted Transcriptome SEQuencing) platform ^12^ (Supplementary Data 1). To identify the cardiomyocytes amongst the other cell types, we first identified the different cell types based on their transcriptomes. k-medoids clustering of the single cell transcriptomes by the RaceID clustering algorithm was used ^13^ (Fig. 1c and Supplementary Data 2), and visualized in two dimensions using *t*-distributed stochastic neighbor embedding (*t*-SNE) (Fig. 1d). A total of 12 cell clusters were identified, including a large group of cardiomyocytes (clusters 1,2,4,7 and 9), a smaller group of endothelial cells (clusters 5,6,8,10 and 12), and some fibroblasts (cluster 3) and immune cells (cluster 11) using the expression of marker genes for specific cell types (Extended Data Fig. S2). Based on the transcriptome clustering, the cardiomyocytes fell into four main transcriptionally-defined clusters (1, 2, 4 and 7), indicating that the injured heart contained subgroups of cardiomyocytes. To address whether the border zone cardiomyocytes were enriched in one of the four cardiomyocyte clusters we compared the mCitrine fluorescence intensity (recorded during FACS sorting) of the cardiomyocyte and found that the average intensity was highest in cluster 7 and lowest in cluster 2 (Extended Data Fig. S4). In addition we analysed the single-cell transcriptome data for the expression of known border zone genes (*nppa, vmhc* and *mustn1b*) and again found that cells expressing these genes were mostly in cluster 7 (Fig. 1e and Extended Data Fig. 3a). Together, these results indicate two things: first, the border zone cardiomyocytes (grouped in cluster 7) can be identified as a separate group in the single-cell RNA-seq data. Secondly, these border zone cardiomyocytes are transcriptionally distinct from remote cardiomyocytes (grouped in cluster 2), while two intermediate cardiomyocyte clusters lie in between.

**Figure 1:**
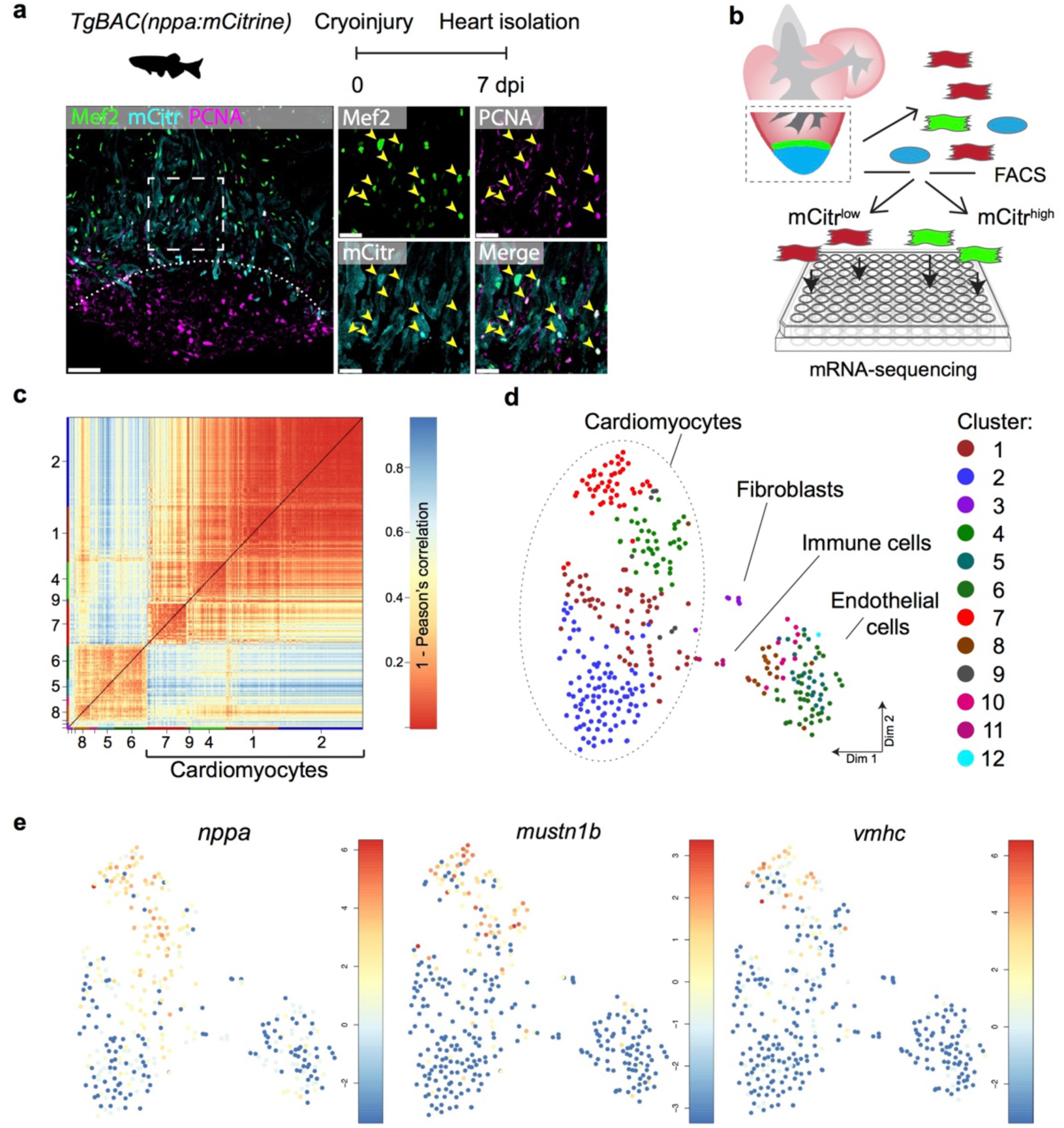
Single-cell mRNA sequencing identifies different cardiomyocyte-populations in the injured zebrafish heart. (a) Schematic of cryoinjury procedure on adult TgBAC(*nppa*:*mCitrine*) fish and immunohistochemistry on section of injured *TgBAC(nppa:mCitrine)* heart 7 days post injury (dpi). Overview image on left and zoom-in of boxed region on the right. Mef2 (in green) labels cardiomyocytes, nppa:mCitrine in cyan, PCNA (in magenta) marks proliferating cells. Arrows indicate triple-positive cells. Dashed line indicates injury site. Scale bar in overview 50 μm. Scale bar in zoom-ins 20 μm. (b) Experimental outline of the single-cell mRNA-sequencing of injured zebrafish hearts (blue, injury area; green, border zone) (c) Pairwise correlation between individual cells across all genes detected. Color-code indicates cell-to-cell distances measured by [1 – Pearson’s correlation coefficient]. StemID clusters are indicated by color and number on the x- and y-axis. (d) t-distributed stochastic neighbor embedding (tSNE) map representation of transcriptome similarities between individual cells. (e)) tSNE maps visualizing log2-transformed read-counts of the border zone marker genes *nppa, mustn1b* and *vmhc*.

Cardiomyocytes in the border zone disassemble sarcomeric structures and re-express markers of embryonic cardiomyocytes suggesting their dedifferentiation ^6–8,14^ We therefore wanted to address the level of dedifferentiation of cluster 7 cardiomyocytes by comparing their transcriptome with embryonic cardiomyocytes. To obtain embryonic cardiomyocytes we performed FACSorting on embryos expressing the cardiomyocyte specific marker *Tg(myl7:GFP)*. Single-cell mRNA-sequencing was performed and combined with the single-cell data from the injured adult hearts (Supplementary Data 3, 4 and 5). The RaceID algorithm identified several cell clusters (Extended Data Fig. S5) with separate clusters for the embryonic and adult cardiomyocytes (Fig. 2a). Importantly, the cluster 7 cardiomyocytes identified in the adult data analysis had a transcriptome that was highly similar to embryonic cardiomyocytes, as shown by pairwise correlation of the differentially expressed genes between the cardiomyocyte clusters: only 184 genes (p-value <0.01), out of 23,707 total detected genes, were differentially expressed between the embryonic and the cluster 7 (border zone) adult cardiomyocytes (Supplementary Data 8). In contrast, over 1800 genes (p-value <0.01) were differentially expressed between embryonic and cluster 2 (remote zone) adult cardiomyocytes.

**Figure 2:**
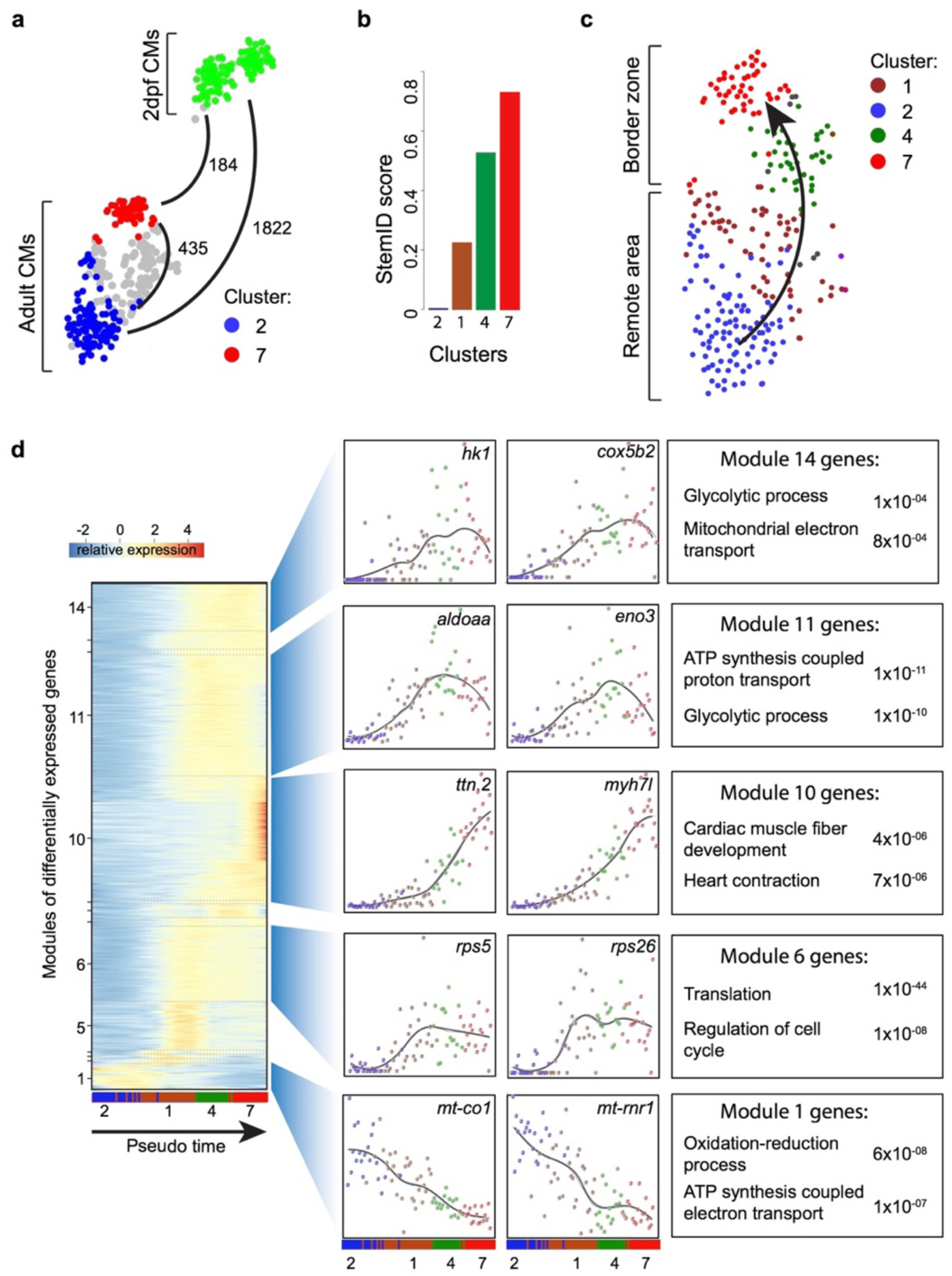
Single-cell analysis reveals dedifferentiation and metabolic changes in border zone cardiomyocytes. (a) tSNE map of adult cardiomyocytes of the injured heart (cluster 7 in red, cluster 2 in blue and clusters 1 and 4 in grey) and embryonic (2 dpf) cardiomyocytes (in green), with number of pairwise differentially expressed genes (p-value<0.01) indicated between cardiomyocyte clusters. (b) Bar plot of StemID scores for the cardiomyocyte clusters (clusters #2, 1, 4 and 7) calculated by the formula: number of significant links for each cluster multiplied by the median transcriptome entropy across all cells in a cluster. (c) Cardiomyocyte clusters from adult injured heart. Arrow indicates the dedifferentiation path derived from the StemID scores. (d) Left ; one-dimensional SOM of z-score transformed expression profiles along the differentiation trajectory incurred by StemID analysis. Y-axis represents the fourteen modules with differentially expressed genes. X-axis represents the pseudo time in which the cells were ordered. Middle; expression profiles of representative genes of the major modules. Y-axis shows transcript counts. X-axis represents the pseudo time. Right; Major gene ontology terms derived from all genes expressed in the module with p-values.

To identify a possible dedifferentiation route we used part of the RaceID algorithm (StemID) that uses the single cell transcriptome data and cell clustering to derive a branched lineage tree ^15^. The algorithm is based on the premise that stem cells and less differentiated cells tend to exhibit more uniform transcriptomes than differentiated cells, which express smaller numbers of genes at higher rates ^16^. Using this approach, we found large differences in transcriptome entropy, resulting in low (cluster 2), intermediate (clusters 1 and 4) and high (cluster 7) StemID scores (Fig. 2b). This gradual increase suggests a dedifferentiation axis from cells in cluster 2 (remote myocardium) to cells in cluster 7 (border zone myocardium) and is in good agreement with our finding that the transcriptome of cluster 7 cardiomyocytes resembles an embryonic cardiomyocyte transcriptome (Fig. 2c). Together, these results indicate that clusters 4 and 7 are enriched for dedifferentiated and proliferative border zone cardiomyocytes while clusters 1 and 2 are enriched for differentiated remote cardiomyocytes.

While cardiomyocytes undergo a well-defined sequence of morphological and transcriptional changes during differentiation, very little is known about the reverse process. Ordering whole-transcriptome profiles of single cells with an unsupervised algorithm can resolve the temporal resolution during differentiation by identifying intermediate stages of differentiation without a priori knowledge of marker genes ^17^. In this manner, the single-cell mRNA-seq experiment will constitute an in-silico time series, with each cell representing a distinct state of differentiation along a continuum. To analyze the transcriptional changes occurring during this apparent dedifferentiation, the most likely dedifferentiation path, based on the StemID scores, was chosen starting at cluster 2 and progressing through clusters 1, 4 and 7. Next, gene expression profiles along this pseudo-temporal order were computed for all detected genes using the single-cell transcriptomes. These gene expression profiles were grouped into modules of co-expressed genes using self-organizing maps (SOMs), resulting in 14 modules (Fig. 2d, Supplementary Data 7). Corroborating our hypothesis of varying differentiation states, we observed that gene expression within these modules changed smoothly over pseudo time. We next analyzed the temporally-ordered expression profiles and identified four groups of genes that shared the same dynamics of expression during this differentiation trajectory. The first group (modules 1, 2) contained genes that were most highly expressed only in cells at the very beginning of the pseudo time line and their expression rapidly declined in cells that were positioned later. This group contained many genes transcribed from mitochondrial DNA, which indicates that the cells at the start of the pseudo time line are mature cardiomyocytes. The second group (module 6) contained genes that are expressed early and stay constant in cells further along the pseudo time line. Many genes involved in translation and cell cycle regulation follow this expression pattern. The third group showed an exponential increase in expression with the highest expression at the end of the pseudo time line (module 10). This group contained genes with a function in cardiac muscle fiber development and heart contraction. The fourth group (modules 11 and 14) contained genes with a low expression in cells both at the start and the end of the pseudo-temporal line and a high expression in cells halfway, suggestive of an early role during the dedifferentiation process. Interestingly this group contained many metabolism genes including glycolytic genes. Together, these data suggest that during dedifferentiation and reentry of the cell cycle, border zone cardiomyocytes undergo profound metabolic changes.

Adult cardiomyocytes rely on fatty acid metabolism and aerobic mitochondrial OXPHOS rather than glycolysis for their energy production ^18^. It was therefore surprising to see that the pseudotime analysis shows a concomitant down regulation of genes transcribed from mitochondrial DNA and upregulation of glycolysis genes. To address mitochondrial activity directly, we measured succinate dehydrogenase (SDH) enzyme activity, located in the inner mitochondrial membrane that functions in both the citric acid cycle and electron transport chain. We observed a 40% reduction in SDH activity specifically in the border zone cardiomyocytes as compared to the remote cardiomyocytes (Fig. 3a). In agreement with the reduced mitochondrial OXPHOS activity, transmission electron microscopy (TEM) imaging revealed more immature mitochondria in border zone cardiomyocytes evidenced by their altered morphology and reduced cristae density (Fig. 3b), which is consistent with previous reports linking mitochondrial function with morphology ^19,20^. Since the pseudotime analysis suggested an upregulation of glycolytic gene expression in the border zone cluster (#7), we performed gene set enrichment analysis (GSEA) for glycolysis genes. The GSEA revealed a strong and significant enrichment in the expression of glycolytic genes in cluster 7 cells compared to cluster 2 cells (Extended Data Fig. S6). We confirmed by in situ hybridization the induced expression in border zone cardiomyocytes of the rate-limiting enzymes hexokinase (*hk1*), pyruvate kinase M1/M2a (*pkma*) and pyruvate dehydrogenase kinase (*pdk2a*), which diverts pyruvate away from the TCA cycle (Fig. 3c and Extended Data Fig.3b). Corroborating the suggested enhanced glycolysis, we observed induced expression of glucose importer genes (*glut1a/slc2a1a* and *glut1b/slc2a1b*) in cluster 7 cells (Extended Data Fig. S6) and enhanced *in vivo* glucose uptake of border zone cardiomyocytes (Extended Data Fig. S7). Furthermore, genes encoding lactate transporters and their proteins were upregulated in cluster 7 and border zone cardiomyocytes (Fig 3d and Extended Data Fig. S6).

**Figure 3:**
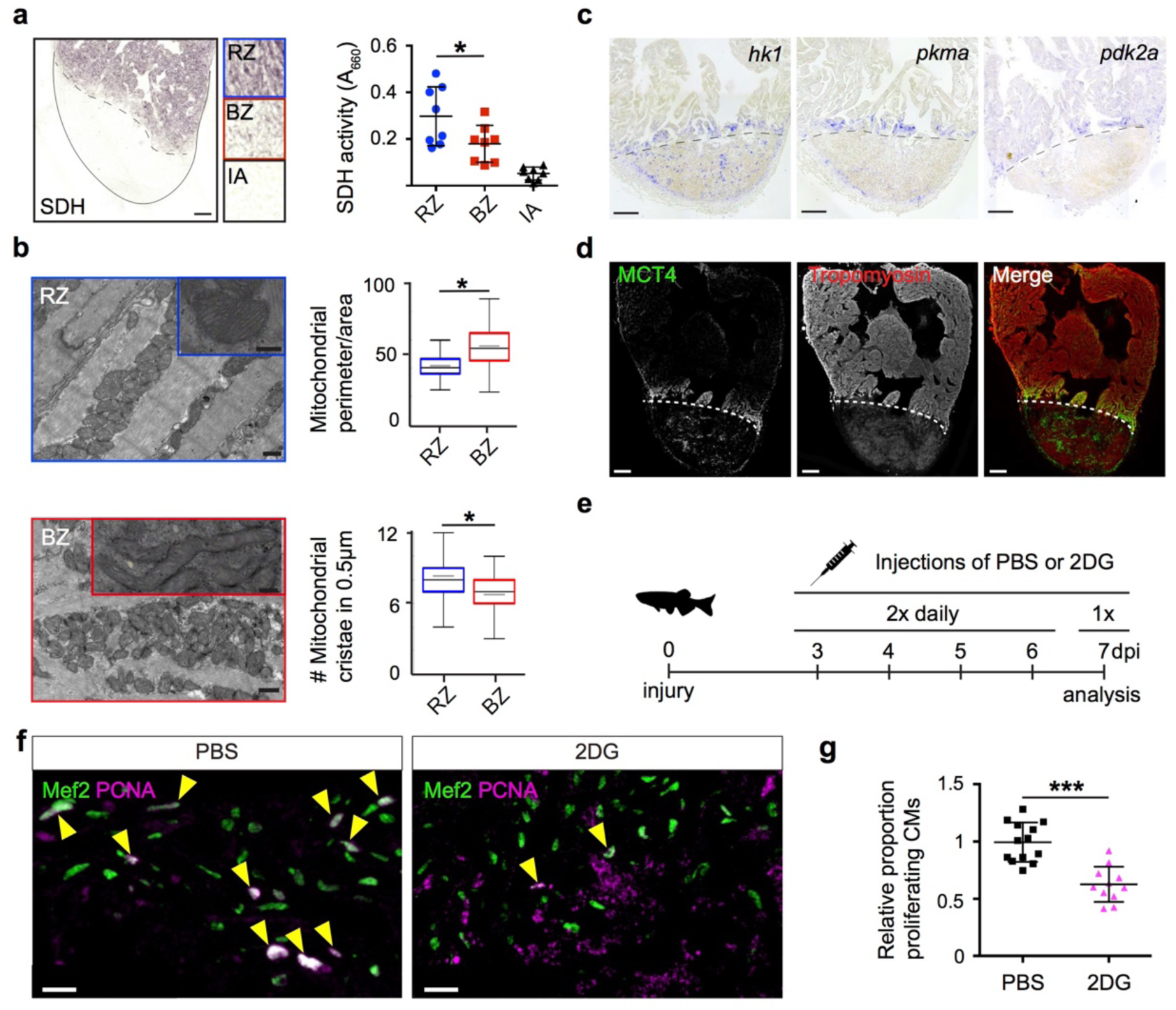
Border zone cardiomyocytes require a metabolic switch from mitochondrial OXPHOS to glycolysis to reenter the cell cycle. (a) Succinate dehydrogenase (SDH) enzyme staining on a 7 dpi heart section with injury area separated by dashed line. Quantification of SDH activity in remote zone (RZ), border zone (BZ) and injury area (IA). (b) Transmission electron microscopy (TEM) images of mitochondria in cardiomyocytes from the remote zone and the border zone of a 7 dpi injured heart. Note the disorganized and irregular shaped mitochondria in the border zone cardiomyocyte. Scale bar 500 nm (200 nm in inserts). Graphs show quantification of mitochondrial perimeter-to-area as a measurement for roundness and quantification of mitochondrial cristae density. * P-value<0.05. (c) In situ hybridizations for glycolytic genes *hk1, pkma* and *pdk2a* on sections of injured zebrafish hearts at 7 dpi. Dashed line indicates injury site. (d) Confocal image of injured zebrafish hearts at 7 dpi stained for the lactate transporter MCT4 (green) and Tropomyosin (red). Dashed line indicates injury site. (e) Experimental design for the 2-DG injections to inhibit glycolysis in injured zebrafish hearts. (f) Confocal image of injured zebrafish hearts at 7 dpi either injected with PBS or 2-DG stained for Mef2c (green) and PCNA (magenta). Arrowheads indicate nuclei positive for Mef2c and PCNA. (g) Quantification of the proliferating cardiomyocytes (double Mef2c/PCNA positive) in the border zone of PBS and 2-DG treated hearts represented as the proportion of proliferating cardiomyocytes compared to the average percentage in the PBS injected group. Each dot represents a single heart (3 sections per heart analyzed). ***, p<0,0001

Together these results suggest a metabolic switch from mitochondrial OXPHOS to glycolysis. To address the functional importance of such a metabolic switch we inhibited glycolysis in injured fish with the glucose analogue 2-Deoxyglucose (2-DG) and analysed its effect on cardiomyocyte proliferation. 2-DG inhibits glycolysis by competitively inhibiting the production of glucose-6-phosphate from glucose. We observed that repeated injections of 2-DG in the adult zebrafish with a cryoinjured heart significantly impaired cardiomyocyte proliferation in the border zone (Fig. 3e-g). Together these observations indicate that, upon cardiac injury, border zone cardiomyocytes switch their metabolism from aerobic mitochondrial OXPHOS to glycolysis, which is necessary for cell cycle reentry.

Next, we investigated the upstream signals that drive the observed metabolic switch in border zone cardiomyocytes during cell cycle reentry. Injury-induced Neuregulin 1 (Nrg1) expression is a potent mitogen that induces cardiomyocyte dedifferentiation and cell cycle reentry by activating ErbB2 receptor signaling ^21^. Furthermore, Nrg1 can induce glucose metabolism in skeletal muscle cells and cardiomyocytes *in vitro* ^22,23^. Therefore, we addressed whether Nrg1 administration can induce glycolysis *in vivo*. To test this we used a previously described transgenic zebrafish model in which Nrg1 overexpression (OE) can be induced in a heart specific manner, *tg(cmlc2:CreER; β-act2:BSNrg1)* ^21^. After tamoxifen injection into *tg(cmlc2:CreER; β-act2:BSNrg1)* fish, we observed a profound and consistent upregulation of glycolysis genes in the myocardium (Fig. 4a). Expression of *hk1*, encoding the rate limiting glycolytic enzyme hexokinase, was strongly induced in the ventricular wall of Nrg1 OE hearts, which correlates well with the observed induction of cardiomyocyte dedifferentiation and proliferation in this region ^21^. From these results, we conclude that glycolysis gene expression can be induced by Nrg1 signaling even in the absence of cardiac injury.

**Figure 4:**
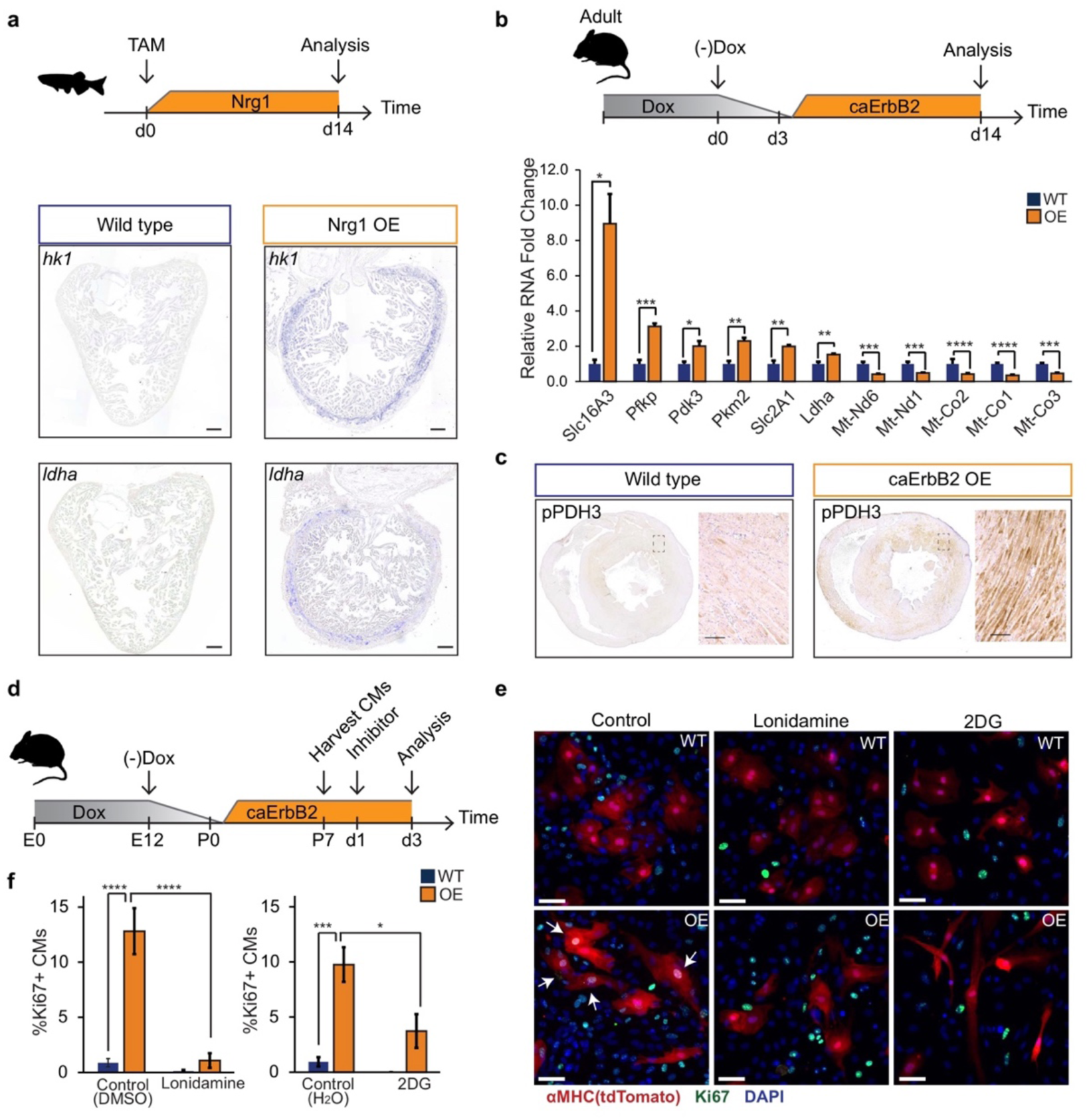
Nrg1/ErbB2 signaling induces proliferation in a glycolysis dependent manner. (a) Cartoon showing experimental procedure to induce cardiomyocyte specific Nrg1 expression in zebrafish. Panels show in situ hybridization for *hexokinase 1* (*hk1*) and *lactate dehydrogenase a* (*ldha*) expression on sections of control hearts (*β-act2:BSNrg1*) and Nrg1 OE hearts (*cmlc2:CreER; β-act2:BSNrg1*), Scale bars represent 100 μm. (b) Upper panel: Cartoon showing the experimental procedure to analyse metabolic gene expression after activating ErbB2 signaling in the murine heart. Lower panel: qPCR results for metabolic genes showing their relative fold change in caErbB2 OE (n=4) heart compared to control WT hearts (n=4). (c) Immunohistochemistry for phospho-PDH3 on sections of control and caErbB2 OE hearts. Scale bar represents 100 μm. (d) Cartoon showing the experimental procedure to analyse the effects of glycolysis inhibitors (2-DG and lonidamine) on cardiomyocyte proliferation. (e) Immunofluorescence analysis on P7 cardiac cultures derived from WT and caErbB2 OE hearts that are endogenously fluorescent for tdTomato under the αMHC promoter, stained for the cell-cycle marker Ki67. Arrows point at Ki67+ CMs. (f) Quantification of % Ki67+ CMs from WT and caErbB2 OE derived fromP7 cardiac cultures treated with the glycolysis inhibitors 2-DG (n=4 for WT and n=4 for OE), and lonidamine (n=7 for WT and n=4 for OE) or their diluents as controls. In all panels, bars represent the mean, and error bars represent s.e.m * p<0.05, ** p<0.01, *** p<0.001, **** p<0.0001, Scale bars represent 50μm

While the mammalian neonatal heart regenerates very efficiently, this regenerative capacity is lost soon after birth ^24,25^. The loss of regenerative capacity in the first week after birth correlates well with the observed change in cardiomyocyte metabolism from glycolysis to mitochondrial OXPHOS ^18^. This has been attributed to an increase in oxygen availability resulting in accumulating reactive oxygen species (ROS) and subsequently DNA damage and reduced proliferation in adult cardiomyocytes ^26–28^. A direct role for glycolysis and its role on cardiomyocyte proliferation and mammalian heart regeneration has not been addressed. To test the hypothesis that glycolysis might be beneficial to cardiomyocyte proliferation and regeneration, we first determined whether a metabolic switch from OXPHOS to glycolysis occurs in a murine model in which regeneration can be induced, such as the transgenic constitutively active ErbB2 receptor overexpression system (caErbB2 OE) ^29^. Consistent with our model, we observed that critical glycolysis genes (e.g. *Pfkp, Pdk3 and Pkm2*) including glucose and lactate transporters (*Slc16A3* and *Slc2A1*) were significantly upregulated in caErbB2 OE cardiomyocytes while genes transcribed from mitochondrial DNA were downregulated (Fig. 4b). *Pdk3* encodes a pyruvate dehydrogenase kinase, which phosphorylates pyruvate dehydrogenase (PDH). PDH is a mitochondrial multi-enzyme complex that converts pyruvate to Acetyl-CoA and provides a primary link between glycolysis and the TCA cycle. Upon phosphorylation by PDK, p-PDH is inactivated and pyruvate is diverted away from the TCA cycle resulting in enhanced lactate production. Consistent with the increase in *pdk3* expression, an increased phosphorylation of PDH was observed in the myocardium of caErbB2 OE hearts (Fig. 4c).

Secondly, we addressed whether the observed switch in metabolic gene expression correlated with enhanced regenerative capacity after injury. Patients and animal models with myocardial infarction (MI) show reduced fatty acid metabolism and increased glucose uptake and glycolysis ^30,31^, which is explained by the reduced oxygen availability in the ischemic region since glycolysis is oxygen independent ^32 33^. Therefore, we performed MI in wild type and caErbB2 OE hearts. Even though glucose uptake and glycolytic enzyme activity in the ischemic area are increased by MI ^30 31^, we observed a stronger upregulation of glycolytic gene expression and decreased mitochondrial gene expression in caErbB2 OE hearts with MI compared to wild type hearts with MI (Extended Data Fig. S8). This stronger and consistent upregulation of glycolytic gene expression in caErbB2 OE hearts correlates with the reported enhanced cardiomyocyte proliferation and improved regeneration ^29^. These findings imply that the enhanced glucose uptake and glycolysis observed after MI might be sufficient for cell survival, but that a stronger induction of glycolysis gene expression is required to fully stimulate cardiomyocyte proliferation and regeneration.

Thirdly, we addressed whether the observed metabolic switch to glycolysis in caErbB2 OE cardiomyocytes is required for their reentry into the cell cycle. Corroborating our model, we indeed observed that treating caErbB2 OE cardiomyocytes *in vitro* with the glycolysis inhibitors 2-DG or lonidamine (inhibiting hexokinase activity) strongly and significantly impaired cell cycle reentry (Fig 4e) and cytokinesis (Extended Data Fig. 9). Together these results demonstrate that in the murine cardiomyocytes, ErbB2 signaling drives a metabolic switch from OXPHOS to glycolysis, which is required for their cell cycle reentry. These results suggest that this metabolic switch in cardiomyocytes is beneficial for heart regeneration.

In conclusion, the use of single cell transcriptomics has allowed us to identify and characterize the proliferating cardiomyocyte population that arises following injury in the regenerating zebrafish heart. Most striking is that these proliferative cardiomyocytes exhibits a metabolic reprogramming hallmark, whereby genes encoding regulators for glucose transport and glycolysis are strongly induced, while simultaneously reducing mitochondrial activity and undergoing dedifferentiation. This metabolic reprogramming phenomenon is necessary for cell cycle reentry of adult cardiomyocytes, and Nrg1/ErbB2 signaling mechanistically acts upstream of this event. More importantly, metabolic reprogramming appears to be a conserved mechanism for cardiomyocyte cell cycle reentry since mammalian models of adult cardiomyocyte proliferation also display this reprogramming hallmark. Likewise, alteration of metabolic states directly influenced proliferative capacity, further demonstrating its crucial role for mammalian cardiomyocyte cell cycle reentry.

This raises the question why cardiomyocytes need to switch their metabolism to enter the cell cycle? Interestingly, a similar metabolic shift from aerobic OXPHOS to glycolysis occurs in proliferating tumor cells, known as the Warburg effect ^34^. Furthermore, progenitor cells in the developing embryo as well as induced pluripotent stem cells, also depend on glycolysis to maintain proliferation and their potency ^35,36^. Glycolysis provides essential metabolites that are needed for cellular reprogramming and proliferation ^37^, but glycolytic enzymes can also directly interact with cell cycle regulators to promote proliferation ^38 39^. Our work suggests that therapeutic targeting of metabolic reprogramming in cardiomyocytes may be beneficial to repair damaged hearts.

## Methods

### Transgenic zebrafish lines and cryoinjury

All procedures involving animals were approved by the local animal experiments committees and performed in compliance with animal welfare laws, guidelines and policies, according to national and European law.

The following fish lines were used: myl7:GFP^twu34Tg^ ^40^. The *Tg(cmlc2:CreER; β-act2:BSNrg1)* line was used as described before ^21^. The *TgBAC*(*nppa*:mCitrine) line was generated essentially as described previously ^41^. In short, an iTOL2_amp cassette for pTarBAC was inserted in the vector sequence of bacterial artificial chromosome (BAC) CH211-70L17, which contains the full *nppa* locus. Subsequently, a mCitrine_kan cassette was inserted at the ATG start codon of the first exon of the nppa gene. Amplification from a pCS2+mCitrine_kanR plasmid was achieved with primers: FWD_NPPA_HA1_GFP 5’-gagccaagccagttcagagggcaagaaaacgcattcagagacactcagagACCATGGTGAGCAAGGGCGAGG-3’ and REV_NPPA_HA2_NEO 5’-gtctgctgccaaaccaggagcagcagtcctgtcagaattagtcccccggcTCAGAAGAACTCGTCAAGAAGGCGATAGAA -3’.

Sequences homologous to the BAC are shown in lower case. Recombineering was performed following the manufacturer’s protocol (Red/ET recombination; Gene Bridges GmbH, Heidelberg, Germany) with minor modifications. BAC DNA isolation was carried out using a Midiprep kit (Life Technologies BV, Bleiswijk, The Netherlands). BAC DNA was injected at a concentration of 300 ng/μl in the presence of 25ng Tol2 mRNA. At 3 dpf, healthy embryos displaying robust nppa-specific fluorescence in the heart were selected and grown to adulthood. Subsequently, founder fish were identified by outcrossing and their progeny grown to adulthood to establish the transgenic line.

Zebrafish of ~ 4 to 18 months of age (males and females, TL strain) were used for regeneration experiments. Cryoinjuries were performed as previously described ^42^, except that a liquid nitrogen-cooled copper filament of 0.3 mm diameter was used instead of dry ice.

Sample sizes were chosen to accommodate the generally accepted standards in the field: 5 or more cryoinjured hearts per condition.

Animals were only excluded from experiments in case of severe sickness/infection/aberrant behavior (according to animal experiment guidelines).

### Transgenic mouse lines and animal procedures

Doxycycline-inducible CM-restricted overexpression of a constitutively active Erbb2 (caErbb2) was generated by crossing the TetRE–caErbb2 ^43^ mouse line with αMHC–tTA which expresses the tetracycline-responsive transcriptional activator (tTA) under the control of the human alpha myosin heavy chain promoter (αMHC) ^44^. Doxycycline (DOX, Harlan Laboratories, TD02503) was administered in the food to repress transgene expression. For cultures derived of OE/WT hearts, we additionally intercrossed the αMHC-cre ROSA26-tdTomato transgenes in order to visualize CMs. For myocardial infarction, mice were sedated with isoflurane (Abbott Laboratories) and were artificially ventilated following tracheal intubation. Experimental myocardial infarction was induced by ligation of the left anterior descending coronary artery (LAD ligation). Following the closure of the thoracic wall mice were warmed for several minutes until recovery.

### Immunofluorescence

#### ADULT

For immunofluorescence, hearts were extracted, fixed in 4% PFA at room temperature for 1,5 h and cryosectioned into 10 μm sections. Heart sections were equally distributed onto seven serial slides so each slide contained sections representing all areas of the ventricle.

Primary antibodies used were anti-AuroraB kinase (BD Transduction laboratories #611082, 1:200), anti-Ki67 (Cell Marque #275R, 1:200), anti-MCT4 (Santa Cruz #SC50329, 1:200), anti-PCNA (Dako #M0879, 1:800), anti-GFP (aves #GFP-1010, 1:1000), and anti-Mef2c (Santa Cruz #SC313, Biorbyt #orb256682 both 1:1000). Antigen retrieval was performed by heating slides containing heart sections at 85°C in 10 mM sodium citrate buffer (pH 6) for 10 minutes. Secondary antibodies conjugated to Alexa 488 (ThermoFisher Scientific), Cy3 or Cy5 (Jackson Laboratories) were used at a dilution of 1:1000. Nuclei were shown by DAPI (4’,6-diamidino-2-phenylindole) staining. Images of immunofluorescence stainings are single optical planes acquired with a Leica Sp8 or Sp5 confocal microscope. Quantifications of PCNA, Mef2, and mCitrine expression were performed in cardiomyocytes situated within 150 μm from the wound border on 3 sections of >4 hearts.

#### EMBRYONIC

Live embryos were immobilized using ms222 and embedded in nitrocellulose + E3 to be mounted on a Leica SPE confocal microscope, followed by a Z-stack maximum projection (step size 2 μm).

#### MAMMALIAN P7 CARDIAC CULTURES

For immunofluorescence, cardiac cultures were fixed with 4% PFA for 10 minutes on room temperature on the shaker, followed by permeabilization with 0.5% Triton X-100 in PBS for 5min, and blocking with 5% bovine serum albumin (BSA) in PBS containing 0.1% Triton for 1h at room temperature.

### Quantitative PCR

RNA from whole hearts was isolated using the MiRNeasy kit (Qiagen, 217004), according to the manufacturer’s instructions. RNA was quantified using a NanoDrop spectrophotometer. A High Capacity cDNA Reverse transcription kit (Applied Biosystems, 4374966) was used to reverse transcribe 1μg of purified RNA according to the manufacturer’s instructions. qPCR reactions were performed using Fast SYBR Green PCR Master Mix (Applied Biosystems). Oligonucleotide sequences for real-time PCR analysis performed in this study are listed in SupplementaryTable 1.

### *In situ* hybridization

#### PARAFFIN

After o/n fixation in 4% PFA, hearts were washed in PBS twice, dehydrated in EtOH, and embedded in paraffin. Serial sections were made at 10 μm. *In situ* hybridization was performed on paraffin-sections as previously described ^45^ except that the hybridization buffer used did not contain heparin and yeast total RNA.

#### CRYOSECTIONS

Sections were obtained as described earlier. *In situ* hybridization was performed as for paraffin, however sections were pre-fixed for 10 minutes in 4% PFA + 0.25% glutaraldehyde before Proteinase K treatment. Moreover, slides were fixed for 1 hour in 4% PFA directly after staining.

### Isolation of single cells from cryoinjured hearts

Cryoinjured hearts (n=13) were extracted at 7 dpi. Cells were dissociated according to^46^. For cell sorting, viable cells were gated by negative DAPI staining and positive YFP-fluorescence. In brief, the FACS gating was adjusted to sort cells for nppa:mCitrine^high^ (to enrich for proliferating cardiomyocytes) and nppa:mCitrine^low^ (remote cardiomyocytes and other cell types) cells. In total n=576 mCitrine^high^ and n=192 mCitrine^low^ cells were sorted into 384-well plates and processed for mRNA sequencing as described below.

### Isolation of single cells from embryonic zebrafish

Transgenic *tg(myl7:GFP)* 2-day-old embryos (n=200) were dechorionated and digested in HBSS Ca^2+^/Mg^2+^ free media containing 0.1% collagenase type II (Gibco) at 32°C for 30–40 minutes followed by 1X TrypLE Express (Gibco) for 15–30 minutes at 32°C with agitation. Dissociated cells were then FACSorted and subjected to single-cell mRNA-seq.

### Single-cell mRNA sequencing

Single-cell sequencing libraries were prepared using SORT-seq ^12^. Live cells were sorted into 384-well plates with Vapor-Lock oil containing a droplet with barcoded primers, spike-in RNA and dNTPs, followed by heat-induced cell lysis and cDNA syntheses using a robotic liquid handler. Primers consisted of a 24 bp polyT stretch, a 4bp random molecular barcode (UMI), a cell-specific 8bp barcode, the 5’ Illumina TruSeq small RNA kit adapter and a T7 promoter. After cell-lysis for 5 minutes at 65°C, RT and second strand mixes were distributed with the Nanodrop II liquid handling platform (Inovadyne). After pooling all cells in one library, the aqueous phase was separated from the oil phase, followed by IVT transcription. The CEL-Seq2 protocol was used for library prep ^47^. Illumina sequencing libraries were prepared with the TruSeq small RNA primers (Illumina) and paired-end sequenced at 75 bp read length on the Illumina NextSeq platform. Mapping was performed against the zebrafish reference assembly version 9 (Zv9).

### Bioinformatic analysis

To analyze the single-cell RNA-seq data we used the previously published RaceID algorithm ^13^. For the adult hearts we had a dataset consisting of two different libraries of 384 cells each for a combined dataset of 768 cells, in which we detected 19257 genes. We detected an average of 10,443 reads per cell. Based on the distribution of the log10 total reads plotted against the frequency, we introduced a cutoff at minimally 3500 reads per cell before further analysis. This reduced the number of cells used in the analysis to 352. Next, we downsampled reads to 3500 unique (UMI corrected) transcripts per cell, as means of normalization. Moreover, we discarded genes that were not detected at >3 transcripts in >1 cell and These cutoffs is a stringent normalization method that allows us to directly compare detected transcripts between cells from different cell types and libraries. Batch-effects were analyzed and showed no plate-specific clustering of certain clusters.

For embryonic cardiomyocytes, two libraries of 384 cells were combined to obtain a set of 768 cells, in which 22213 genes could be detected. An average of 4455 reads per cell was detected. After downsampling of this library to 3500 reads per cell 312 cells were included for further analysis. Further analysis was performed by combining the embryonic heart data with the injured adult heart data (total of 664 cells).

The StemID algorithm were used as previously published ^15^. In short, StemID is an approach developed for inferring the existence of stem cell populations from single-cell transcriptomics data. StemID calculates all pairwise cell-to-cell distances (1 – Pearson correlation) and uses this to cluster similar cells into clusters that correspond to the cell types present in the tissue. The StemID algorithm calculates the number of links between clusters. This is based on the assumption that cell types with less links are more canalized while cell types with a higher number of links have a higher diversity of cell fates. Besides the number of links, the StemID algorithm also calculates the change in transcriptome entropy. Differentiated cells usually express a small number of genes at high levels in order to perform cell specific functions, which is reflected by a low entropy. Stem cells and progenitor cells display a more diverse transcriptome reflected by high entropy ^16^. By calculating the number of links of one cluster to other clusters and multiplying this with the change in entropy, it generates a StemID score, which is representative to “stemness” of a cell population.

### Inference of co-expressed gene modules

To identify modules of co-expressed genes along a specific differentiation trajectory (defined as a succession of significant links between clusters as identified by StemID) all cells assigned to these links were assembled in pseudo-temporal order based on their projection coordinate. Next, all genes that are not present with at least two transcripts in at least a single cell are discarded from the sub-sequent analysis. Subsequently, a local regression of the z-transformed expression profile for each gene is computed along the differentiation trajectory. These pseudo-temporal gene expression profiles are topologically ordered by computing a one-dimensional self-organizing map (SOM) with 1,000 nodes. Due to the large number of nodes relative to the number of clustered profiles, similar profiles are assigned to the same node. Only nodes with more than 3 assigned profiles are retained for visualization of co-expressed gene modules. Neighboring nodes with average profiles exhibiting a Pearson’s correlation coefficient >0.9 are merged to common gene expression modules. These modules are depicted in the final map. Analyses were performed as previously published ^15^.

### Transmission Electron Microscopy

Hearts were excised and immediately chemically fixated at room temperature with 2,5% glutaraldehyde and 2% formaldehyde (EMS, Hainfield USA) in 0.1M phosphate buffer pH 7.4 for 2 hr. Next, hearts were post fixed with 1 % OsO_4_ (EMS, Hainfield USA)/ 1.5 % K_3_Fe(CN)_6_ in 0.065 M phosphate buffer for 2 h at 4 °C and finally 1 h with 0,5% uranyl acetate. After fixation, hearts were dehydrated in a graded series of acetone and embedded in Epon epoxy resin (Polysciences). Ultrathin sections of 60 nm were cut on a Leica Ultracut T (Leica, Vienna, Austria) and contrasted with uranyl acetate (0.4% in AD, EMS, Hainfield USA) and lead citrate (Leica Vienna, Austria) using the AC20 (Leica Vienna, Austria) and examined with a Jeol 1010 electron microscope (Jeol Europe, Nieuw Vennep, The Netherlands).

### Quantification of mitochondrial parameters

In every heart, each in the borderzone and remote myocardial region, 100 well-deliniated mitochondria with clearly visible outer and inner membranes were selected. Mitochondrial perimeter and surface were measured using the freehand tool of Image J. The perimeter to surface ratio was calculated and used as a factor that describes the pluriformity of mitochondria. The amount of cristae was estimated by counting the number of cristae intersected by a line of 0.5μm length in 40 mitochondria per region.

### Histology and enzyme histochemistry

Serial cryosections of the heart were cut 7μm thick and either fixed in formalin, stained with Meyer’s hematoxylin and eosin (HE), dehydrated and mounted in Entellan, or incubated for enzyme histochemistry. Chemicals for histochemistry of succinate dehydrogenase (SDH) activity were obtained from Sigma Aldrich. Sections for SDH activity were incubated for 20 min at 28°C in 37.5 mM sodium phosphate buffer pH 7.60, 70 mM sodium succinate, 5 mM sodium azide and 0.4 mM tetranitro blue tetrazolium (TNBT). The reaction was stopped in 10mM HCl. Controls without succinate did not stain. The incubated sections were mounted in glycerine gelatin. The absorbances of the SDH-reaction product in the sections were determined at 660 nm using a calibrated microdensitometer and ImageJ.

### Pharmacological inhibition of glycolysis

#### ZEBRAFISH

Zebrafish were injured and received intraperitoneal (i.p.) injections twice daily with either PBS or 2-Deoxy-D-Glucose (Sigma-Aldrich, 1mg/g) from days 3 to 6 and one more injection on day 7 after injury, two hours before fish were euthanized and hearts harvested. I.p. injections were performed using a Hamilton Syringe (gauge 30) as described in literature ^48^. Injection volumes were corrected to body weight (30μl/g).

#### MAMMALIAN P7 CARDIAC CULTURES

Primary cardiac cultures were isolated from P7 mice using a neonatal dissociation kit (Miltenyi Biotec,130-098-373) using the gentleMACS homogenizer, according to the manufacturer’s instructions and cultured in Gelatin-coated (0.1%, G1393, Sigma) wells with DMEM/F12 medium supplemented with L-glutamine, Na-pyruvate, nonessential amino acids, penicillin, streptomycin, 5% horse serum and 10% FBS (‘complete-medium’) at 37°C and 5% CO2 for 24h. Afterwards, medium was replaced with FBS-depleted medium (otherwise same composition) for additional 48hours of culture in either 3mM 2DG (Sigma-Aldrich) or 80μΜ lonidamine before further processing.

### *Ex vivo* glucose uptake

Fish were euthanized on ice water before hearts were extracted in PBS + heparin and were allowed to bleed out for 15 minutes. Hearts were then transferred into fresh PBS + 10%KCl to stop the heart from beating and mounted directly on a glass bottom cell culture dish in 1% agarose. Thereafter, 2NBDG (Caymanchem #11046, 400μM) was added to the dish and the hearts were taken directly for imaging. Imaging was performed using a Leica SP5 multiphoton microscopy using 930nm laser excitation wavelength. 150μm z-stacks were made with z-step size 5μm every 5 minutes for 2 hours.

### Gene set enrichment analysis (GSEA)

GSEA (Genepattern, Broad Institute) was performed to assess enrichment for glycolytic genes upregulated between cluster 7 and 2. A list of genes involved in zebrafish glycolysis was obtained from KEGG. As number of permutations 1000 was used, which means p=0 indicates p<0.001.

## Acknowledgements

We would like to thank V. Christoffels for critical reading of the manuscript and Life Science Editors for editing support. J.B. acknowledges support from the Netherlands Cardiovascular Research Initiative and the Dutch Heart Foundation (grants CVON2011-12 HUSTCARE and Cobra^3^) and ERA-CVD grant CARDIO-PRO JCT2016-40-080. L.G. was supported by an EMBO long-term fellowship.

## Author contributions

J.B. and E.T. conceived and designed the project. F.T. and J.C.P. generated the nppa-reporter line. F.K., D.E.M.d.B, P.N., M.J.M., D.G. and A.v.O. performed the single cell sequencing and helped with bioinformatics analyses. L.G. performed embryonic zebrafish work. D.E.M.d.B., H.H. and F.K. performed cryoinjuries, immunohistochemical analysis and in situ hybridizations. C.d.H, G.P. and J.K. performed and analyzed the electron microscopy data. W.N., W.J.v.d.L. and R.T.J. performed and analyzed enzymatic stainings. A.A. and A.S. performed and analysed mouse experiments. J.B., H.H., D.E.M.d.B, F.K and P.N. all helped writing the manuscript. All authors approved the manuscript.

---

**Supplemental Figure 1:**
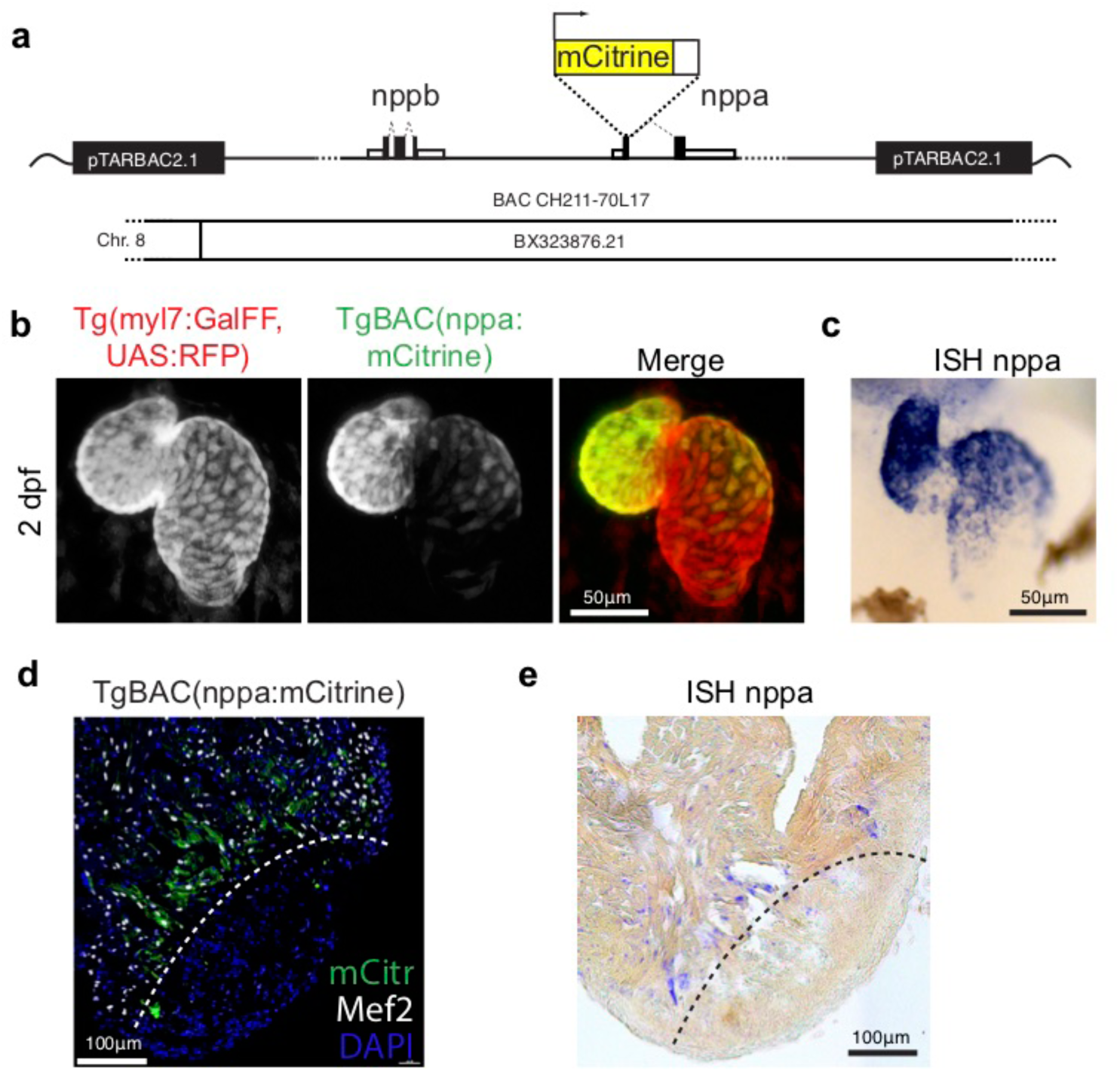
*TgBAC(nppa:mCitrine)* expression recapitulates endogenous *nppa* gene expression. (a) Design of the bacterial artificial chromosome (BAC) used for generation of the transgenic line TgBAC(*nppa*:mCitrine). (b) Transgenic mCitrine expression in the heart in relation to RFP expression in the whole myocardium of Tg(*myl7*:GFF, *UAS*:RFP, *nppa*:mCitrine) in embryos at 2-days post-fertilization (dpf). (c) Whole mount *in situ* hybridization for endogenous *nppa* expression in embryos at 2 dpf. Note the specific expression in ventricle and atrium and absence of expression in atrioventricular canal of the *nppa:mCitrine* transgene expression (in panel b) as well as for the endogenous *nppa* expression (in panel c). (d, e) nppa:mCitrine expression (d) and endogenous *nppa* expression (e) in the adult heart at 7 days post-injury (dpi). Dotted line indicates the border zone.

**Supplemental Figure 2:**
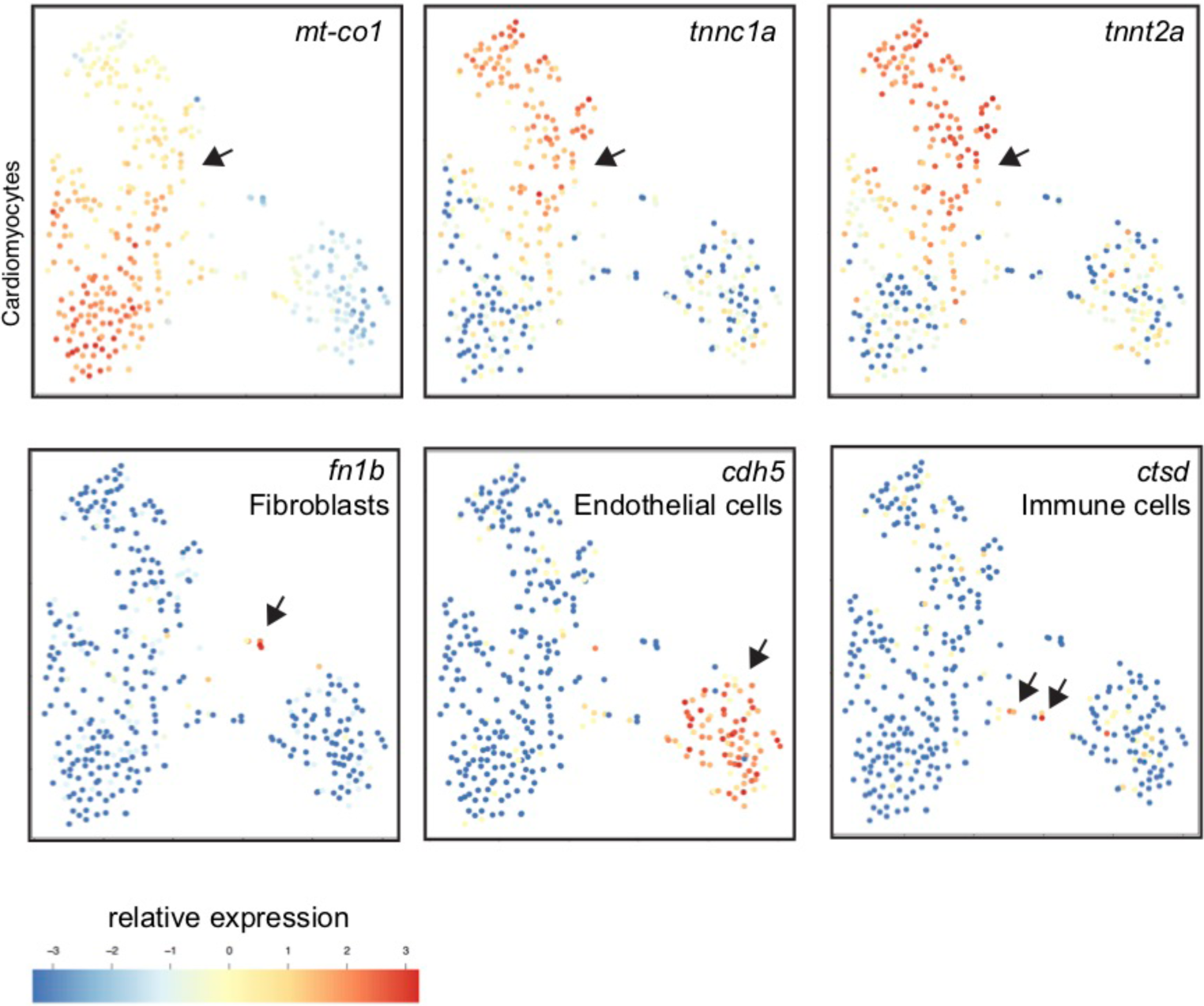
Single-cell mRNA sequencing identifies different cell-populations in the injured zebrafish heart. tSNE maps visualizing log2-transformed read-counts of genes with high expression in cardiomyocytes (*mt-co1*, *tnnc1a, tnnt2a*), in fibroblasts (*fn1b*), in endothelial cells (*cdh5*) and immune cells (*ctsd*). Arrow indicates positive cell populations.

**Supplemental Figure 3:**
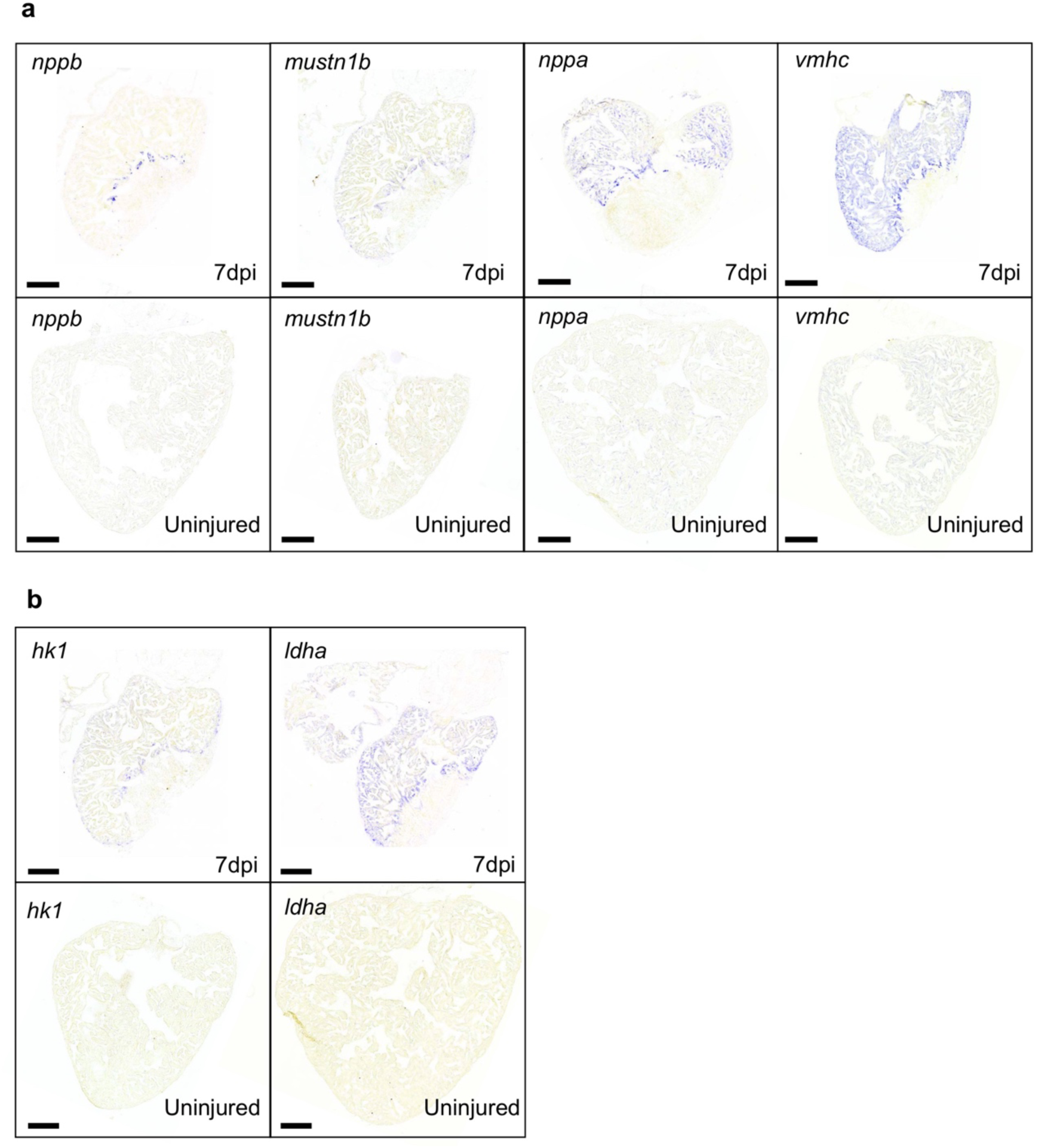
Cardiomyocyte populations with enhanced expression of genes elevated in cluster 7 versus cluster 2 are absent in the uninjured heart. (a) *In situ* hybridizations for border zone genes *nppb, mustn1, nppa* and *vmhc* in zebrafish hearts at 7 dpi compared to uninjured hearts. (b) *In situ* hybridizations for glycolysis genes *hk1* and *ldha* in zebrafish hearts at 7 dpi compared to uninjured hearts. Staining for injured and uninjured hearts was stopped simultaneously. Scale bars indicate 200μm. (n=3 per condition)

**Supplemental Figure 4:**
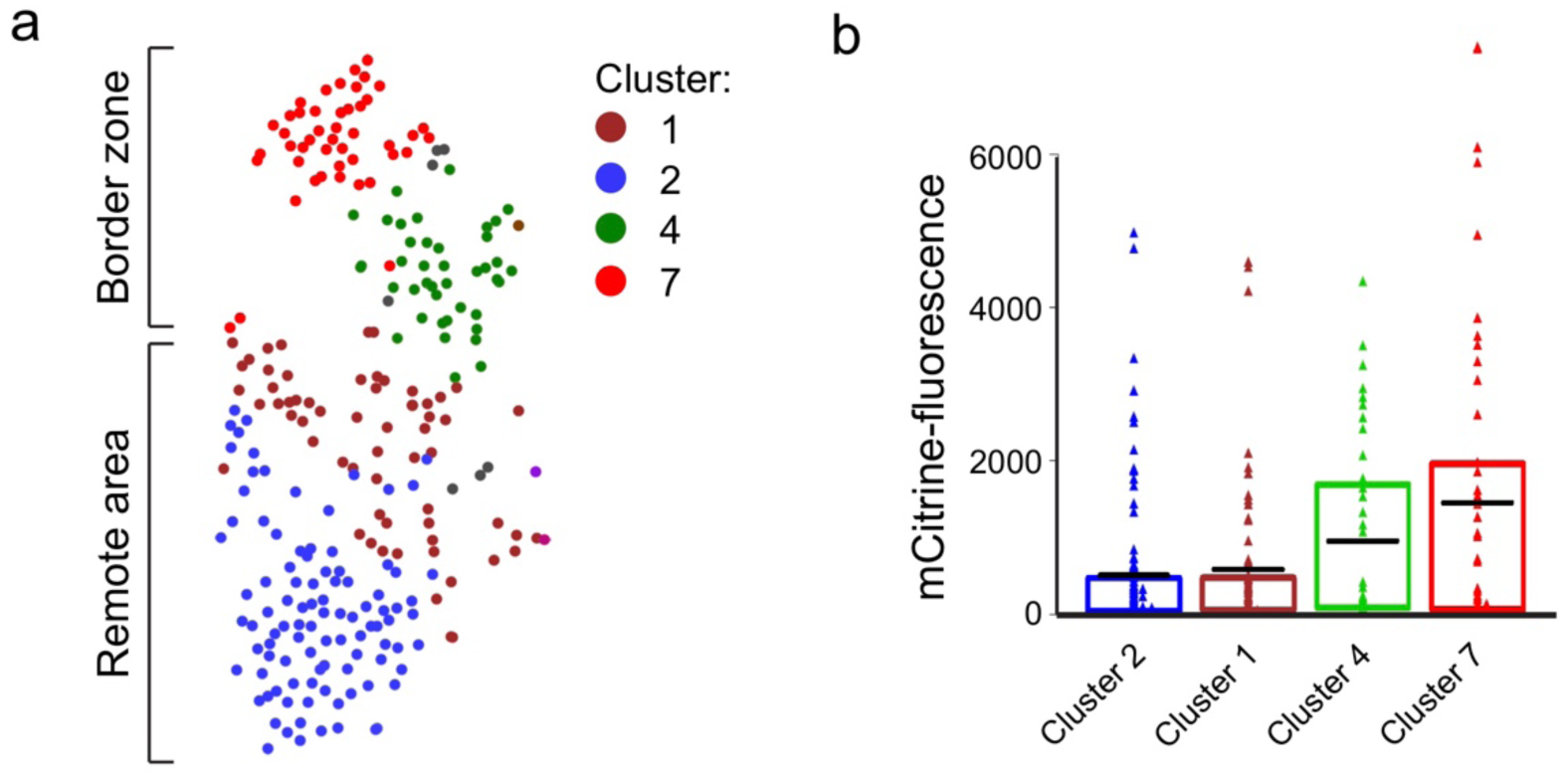
Cluster 7 cells display highest nppa:mCitrine expression. (a) tSNE map of the cardiomyocyte clusters derived from the single-cell mRNA-sequencing. (b) mCitrine fluorescence levels of the nppa:mCitrine FACS sorted cells from injured hearts that were used for the single-cell RNA-sequencing. Triangles represent individual cells of the four cardiomyocyte clusters. The box indicates the 25–75% quartiles, black lines indicate mean-fluorescence per cluster.

**Supplemental Figure 5:**
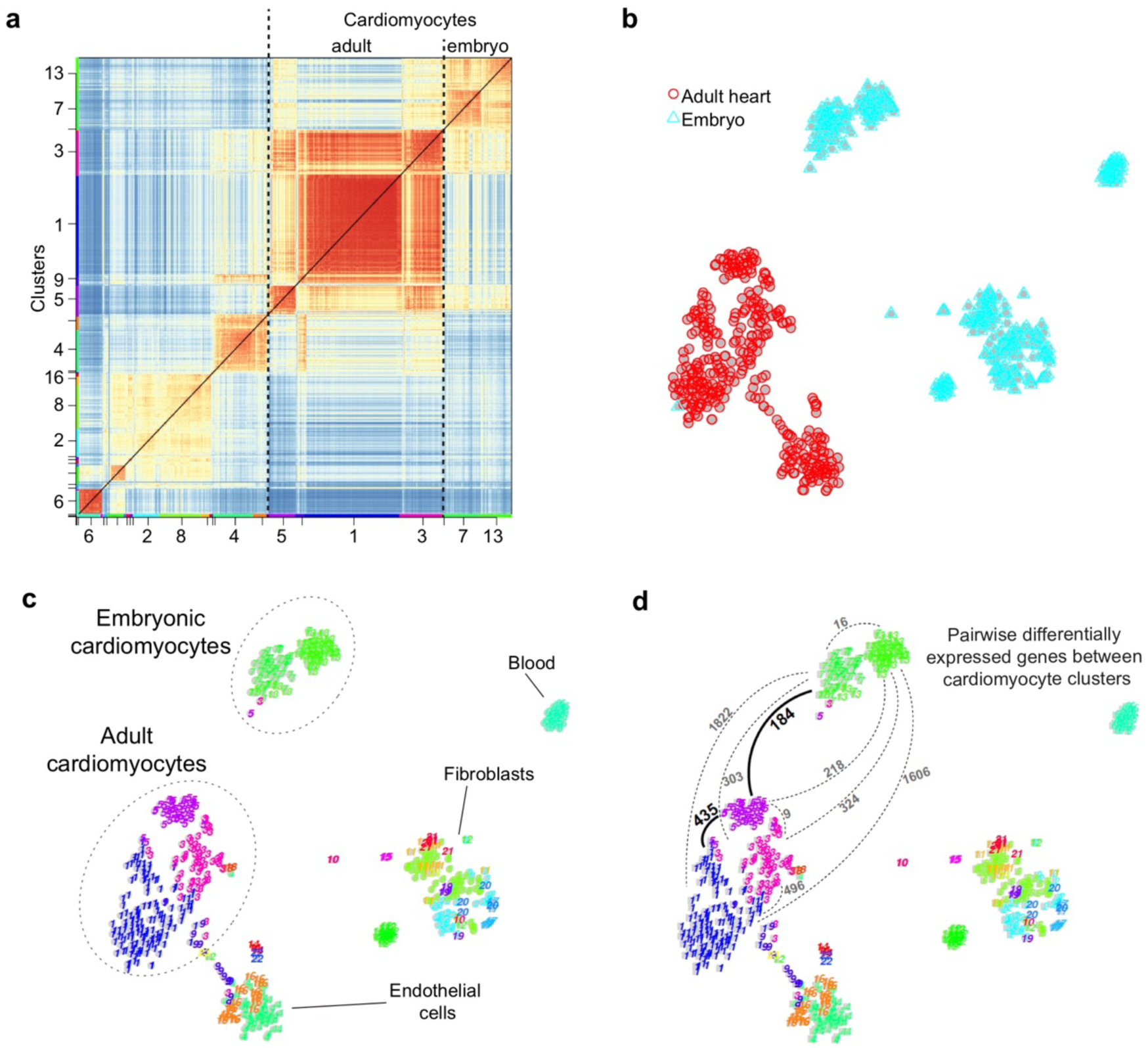
Border zone cardiomyocytes resemble embryonic cardiomyocytes. (a) Pairwise correlation between individual cells derived from the adult heart and whole embryo FACS sorts across all genes. Color-code indicates cell-to-cell distances measured by [1 – Pearson’s correlation coefficient]. StemID clusters are indicated on the x- and y-axis. Adult cardiomyocyte clusters are colored in blue, pink and magenta. Embryonic cardiomyocyte clusters are colored in green (b) tSNE map showing the origin (adult or embryonic) of the clustered cells. Injured adult heart in red and 2 dpf embryonic heart in cyan. (c) tSNE map representation of transcriptome similarities between individual cells and the identified clusters. 2-dimensional arrangement of the clusters identified by StemID and the suggested cell-type associated with each cluster are shown. (d) tSNE map of the clustered cells, with number of pairwise differentially expressed genes (p-value<0.01) indicated between cardiomyocyte clusters. Related to extended data 4.

**Supplemental Figure 6:**
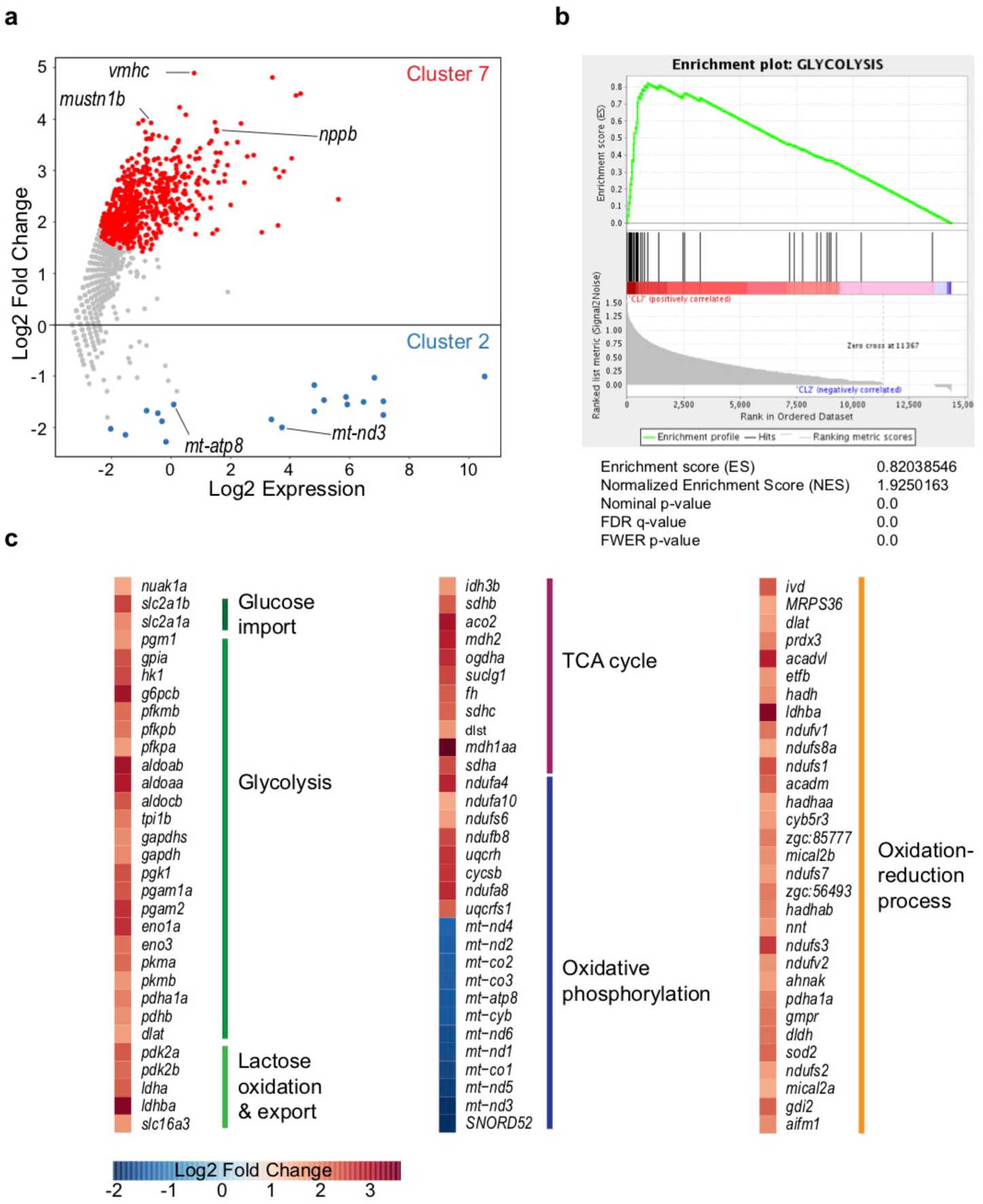
Energy metabolism genes are differentially expressed between cluster 7 and cluster 2 cells. (a) Plot showing differentially expressed genes between cluster 7 versus cluster 2 cells. Differentially expressed genes (p-value<0.05) are highlighted in red (upregulated in cluster 7 cells) and blue (upregulated in cluster 2 cells). Complete gene list can be found in Supplementary Data8. 771 genes were differentially expressed (p < 0.05; adjusted p-value after Benjamini-Hochberg correction), of which 752 were specifically upregulated in cluster 7 cardiomyocytes, including the border zone genes, *nppa*, *nppb* and *mustn1b*. (b) Gene set enrichment analysis for glycolysis genes on all genes with differential expression between cluster 7 versus cluster 2 cells (p<0.05). (c) Log2 fold change of differentially expressed genes (cluster 7 vs cluster 2) with a function in energy metabolism. High relative expression in cluster 7 depicted in red, high expression in cluster 2 depicted in blue.

**Supplemental Figure 7:**
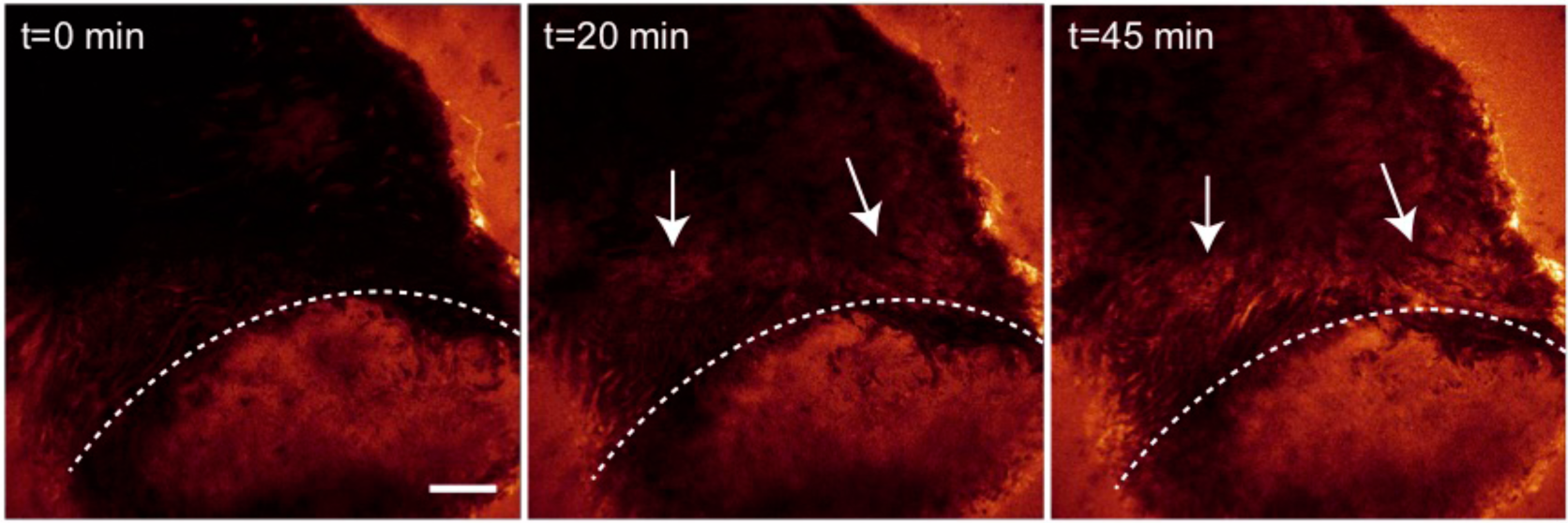
Enhanced glucose uptake by border zone cells. Time-lapse multi-photon confocal images of whole heart. The heart was isolated 7 days post injury and incubated with 2-NBDG, a fluorescent glucose analogue, at t=0. Confocal images were taken every 5 min to monitor cellular glucose uptake. Dotted line indicates injury area. Arrows point to regions of the border zone where 2-NBDG uptake is enhanced compared to the remote area. Scale bar represents 100 μm.

**Supplemental Figure 8:**
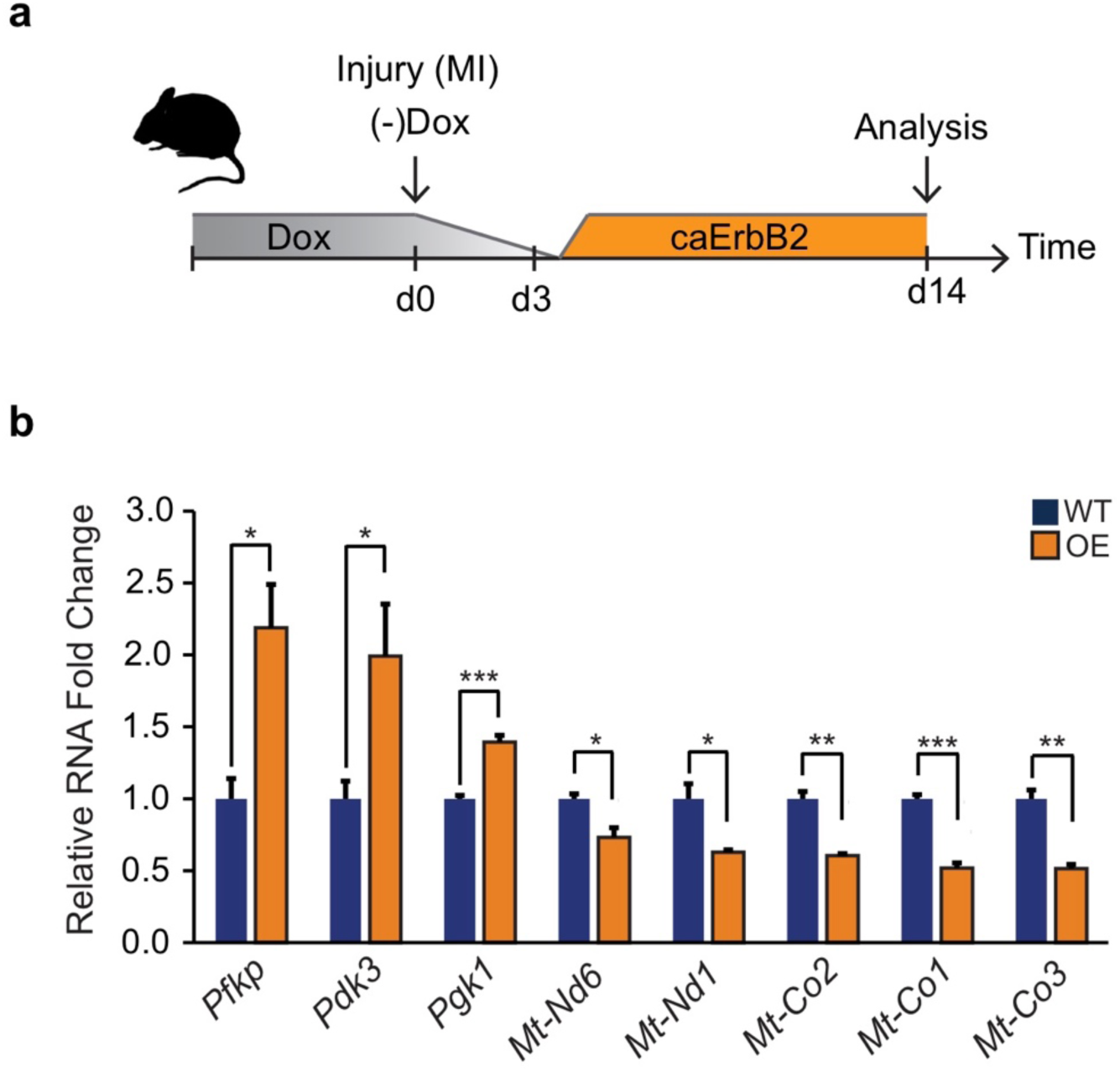
Activation of ErbB2 signaling in murine MI model induces glycolytic gene expression while repressing mitochondrial genes. (a) Cartoon showing experimental procedure. Myocardial infarction (MI) was induced by left anterior descending coronary artery ligation at the same day when Dox was removed to induce cardiomyocyte specific caErbB2 overexpression (OE). Whole hearts where isolated for mRNA extraction 14 days after the MI and caErbB2 induction. (b) qPCR results with relative mRNA fold changes comparing wild type (WT, blue bars) with caErbB2 OE (OE, orange bars). Note the significant upregulation of glycolytic genes (*Pfkp, Pdk3* and *Pgk1*) and significant downregulation of genes transcribed from the mitochondrial DNA (*Mt-Nd1, Mt-Co2, Mt-Co1* and *Mt-Co3*). * p<0,05; ** p<0,01; *** p<0,001

**Supplemental Figure 9:**
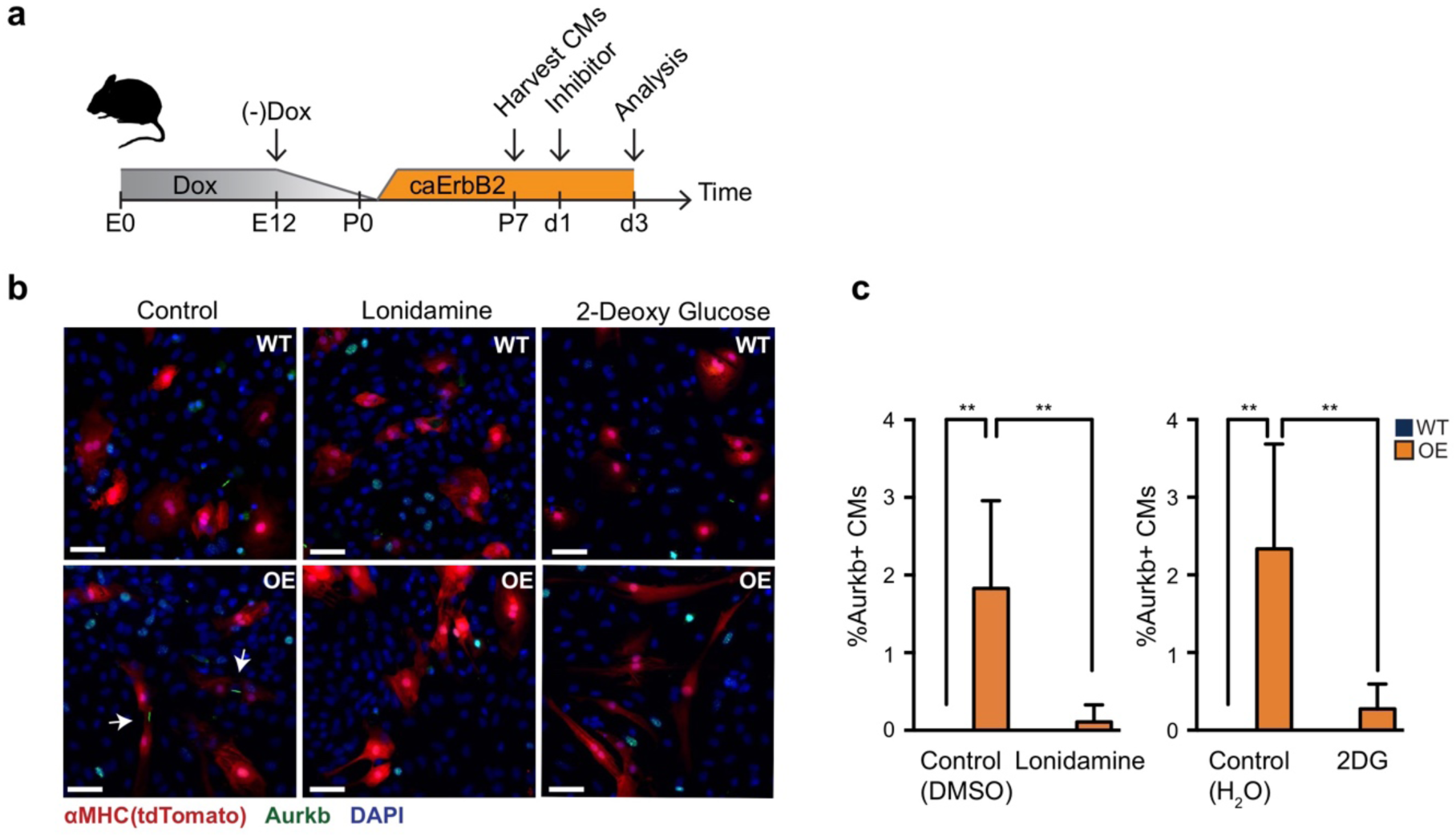
Glycolysis is required for caErbB2 induced cell division. (a) Cartoon showing the experimental procedure to analyse the effects of glycolysis inhibitors (2-DG and lonidamine) on cardiomyocyte proliferation. (e) Confocal images of cells from control and caErbB2 OE hearts stained for the cytokinesis marker AuroraB kinase (green) and myosin heavy chain (aMHC, red) to identify cardiomyocytes and DAPI (Blue). Cells were either control treated or treated with the glycolysis inhibitors Lonidamine or 2-DG. Arrows point to dividing cardiomyocytes. Scale bar represents 50 μm. ** p<0,01

## References

1. Bergmann, O. et al. Dynamics of Cell Generation and Turnover in the Human Heart. Cell 161, 1566–1575 (2015).

2. Bergmann, O. et al. Evidence for Cardiomyocyte Renewal in Humans. Science 324, 98–102 (2009).

3. Senyo, S. E. et al. Mammalian heart renewal by pre-existing cardiomyocytes. Nature 493, 433–436 (2012).

4. Becker, R. O., Chapin, S. & Sherry, R. Regeneration of the ventricular myocardium in amphibians. Nature 248, 145–147 (1974).

5. Poss, K. D., Wilson, L. G. & Keating, M. T. Heart regeneration in zebrafish. Science 298, 2188–2190 (2002).

6. Kikuchi, K. et al. Primary contribution to zebrafish heart regeneration by gata4. Nature 464, 601–605 (2010).

7. Jopling, C. et al. Zebrafish heart regeneration occurs by cardiomyocyte dedifferentiation and proliferation. Nature 464, 606–609 (2010).

8. Wu, C.-C. et al. Spatially Resolved Genome-wide Transcriptional Profiling Identifies BMP Signaling as Essential Regulator of Zebrafish Cardiomyocyte Regeneration. Dev Cell 36, 36–49 (2016).

9. Kang, J. et al. Modulation of tissue repair by regeneration enhancer elements. Nature 532, 201–206 (2016).

10. Lien, C.-L., Schebesta, M., Makino, S., Weber, G. J. & Keating, M. T. Gene expression analysis of zebrafish heart regeneration. PLoS Biol 4, e260 (2006).

11. Sleep, E. et al. Transcriptomics approach to investigate zebrafish heart regeneration. Journal of Cardiovascular Medicine 11, 369–380 (2010).

12. Muraro, M. J. et al. A Single-Cell Transcriptome Atlas of the Human Pancreas. Cell Systems 3, 385–394.e3 (2016).

13. Grün, D. et al. Single-cell messenger RNA sequencing reveals rare intestinal cell types. Nature 525, 251–255 (2015).

14. Lepilina, A. et al. A dynamic epicardial injury response supports progenitor cell activity during zebrafish heart regeneration. Cell 127, 607–619 (2006).

15. Grün, D. et al. De Novo Prediction of Stem Cell Identity using Single-Cell Transcriptome Data. Cell Stem Cell 19, 266–277 (2016).

16. Banerji, C. R. S. et al. Cellular network entropy as the energy potential in Waddington’s differentiation landscape. Sci. Rep. 3, 1129–7 (2013).

17. Trapnell, C. et al. The dynamics and regulators of cell fate decisions are revealed by pseudotemporal ordering of single cells. Nat Biotechnol 32, 381–386 (2014).

18. Lopaschuk, G. D., Collins-Nakai, R. L. & Itoi, T. Developmental changes in energy substrate use by the heart. Cardiovascular Research 26, 1172–1180 (1992).

19. Giraud, M.-F. et al. Is there a relationship between the supramolecular organization of the mitochondrial ATP synthase and the formation of cristae? Biochim. Biophys. Acta 1555, 174–180 (2002).

20. Paumard, P. et al. The ATP synthase is involved in generating mitochondrial cristae morphology. EMBO J 21, 221–230 (2002).

21. Gemberling, M., Karra, R., Dickson, A. L. & Poss, K. D. Nrg1 is an injury-induced cardiomyocyte mitogen for the endogenous heart regeneration program in zebrafish. eLife Sciences 4, (2015).

22. Suárez, E. et al. A Novel Role of Neuregulin in Skeletal Muscle. J Biol Chem 276, 18257–18264 (2001).

23. Cote, G. M., Miller, T. A., LeBrasseur, N. K., Kuramochi, Y. & Sawyer, D. B. Neuregulin-1α and β isoform expression in cardiac microvascular endothelial cells and function in cardiac myocytes in vitro. Exp Cell Res 311, 135–146 (2005).

24. Haubner, B. J. et al. Functional Recovery of a Human Neonatal Heart After Severe Myocardial Infarction. Circ Res 118, 216–221 (2016).

25. Porrello, E. R. et al. Transient regenerative potential of the neonatal mouse heart. Science 331, 1078–1080 (2011).

26. Nakada, Y. et al. Hypoxia induces heart regeneration in adult mice. Nature 541, 222–227 (2016).

27. Puente, B. N. et al. The Oxygen-Rich Postnatal Environment Induces Cardiomyocyte Cell-Cycle Arrest through DNA Damage Response. Cell 157, 565–579 (2014).

28. Tao, G. et al. Pitx2 promotes heart repair by activating the antioxidant response after cardiac injury. Nature 534, 119–123 (2016).

29. D’Uva, G. et al. ERBB2 triggers mammalian heart regeneration by promoting cardiomyocyte dedifferentiation and proliferation. Nat Cell Biol 17, 627–638 (2015).

30. Schelbert, H. R. & Buxton, D. Insights into coronary artery disease gained from metabolic imaging. Circulation 78, 496–505 (1988).

31. Owen, P., Thomas, M. & Opie, L. Relative changes in free-fatty-acid and glucose utilisation by ischaemic myocardium after coronary-artery occlusion. Lancet 1, 1187–1190 (1969).

32. Opie, L. H. Acute metabolic response in myocardial infarction. Heart 33, 129–137 (1971).

33. Scheuer, J. Myocardial metabolism in cardiac hypoxia. Am J Cardiol 19, 385–392 (1967).

34. Warburg, O., Wind, F. & Negelein, E. The metabolism of tumors in the body. J. Gen. Physiol. 8, 519–530 (1927).

35. Folmes, C. D. L. et al. Short Article. Cell Metabolism 14, 264–271 (2011).

36. Gu, W. et al. Glycolytic Metabolism Plays a Functional Role in Regulating Human Pluripotent Stem Cell State. Stem Cell 19, 476–490 (2016).

37. Vander Heiden, M. G., Cantley, L. C. & Thompson, C. B. Understanding the Warburg effect: the metabolic requirements of cell proliferation. Science 324, 1029–1033 (2009).

38. Yang, W. et al. Nuclear PKM2 regulates β-catenin transactivation upon EGFR activation. Nature 480, 118–122 (2011).

39. Dasgupta, S. et al. Metabolic enzyme PFKFB4 activates transcriptional coactivator SRC-3 to drive breast cancer. Nature 556, 249–254 (2018).

40. Huang, C.-J., Tu, C.-T., Hsiao, C.-D., Hsieh, F.-J. & Tsai, H.-J. Germ-line transmission of a myocardium-specific GFP transgene reveals critical regulatory elements in the cardiac myosin light chain 2 promoter of zebrafish. Dev Dyn 228, 30–40 (2003).

41. Bussmann, J. & Schulte-Merker, S. Rapid BAC selection for tol2-mediated transgenesis in zebrafish. Development 138, 4327–4332 (2011).

42. Schnabel, K., Wu, C.-C., Kurth, T. & Weidinger, G. Regeneration of Cryoinjury Induced Necrotic Heart Lesions in Zebrafish Is Associated with Epicardial Activation and Cardiomyocyte Proliferation. PLoS ONE 6, e18503 (2011).

43. Xie, W., Chow, L. T., Paterson, A. J., Chin, E. & Kudlow, J. E. Conditional expression of the ErbB2 oncogene elicits reversible hyperplasia in stratified epithelia and up-regulation of TGFalpha expression in transgenic mice. Oncogene 18, 3593–3607 (1999).

44. Yu, Z., Redfern, C. S. & Fishman, G. I. Conditional transgene expression in the heart. Circ Res 79, 691–697 (1996).

45. Moorman, A., Houweling, A., de Boer, P. & Christoffels, V. Sensitive nonradioactive detection of mRNA in tissue sections: novel application of the whole-mount in situ hybridization protocol. Journal of Histochemistry & Cytochemistry 49, 1 (2001).

46. Tessadori, F. et al. Identification and Functional Characterization of Cardiac Pacemaker Cells in Zebrafish. PLoS ONE 7, e47644 (2012).

47. Hashimshony, T. et al. CEL-Seq2: sensitive highly-multiplexed single-cell RNA-Seq. Genome biology 17, 1–7 (2016).

48. Kinkel, M. D., Eames, S. C., Philipson, L. H. & Prince, V. E. Intraperitoneal injection into adult zebrafish. J Vis Exp (2010). doi:10.3791/2126

